# Evolutionary genomics of host-transposon conflict, multilevel selection, and Red Queen dynamics

**DOI:** 10.64898/2026.07.01.735732

**Authors:** Manaswini Parija, Spandan Patra, Neelesh Dahanukar

**Affiliations:** Department of Life Sciences, School of Natural Sciences, Shiv Nadar Institution of Eminence, Delhi-NCR, India

**Keywords:** Transposable elements, Transposon tolerance, Co-evolutionary arms race, Ancestral trait reconstruction

## Abstract

Transposable elements (TE) jump from one genomic locus to another. Since increase in their copy number is a metabolic burden for the host, TE are considered as genomic parasites. Although host-TE co-existence is regarded as an evolutionary arms race, the hypothesis is not extensively tested especially using evolutionary genomics. We provide a hypothesis testing framework to understand the distribution of TE in genic regions of the host genome, variation in the regulation of TE by host, and effect of these two factors on host-TE co-evolutionary dynamics. We test our hypothesis by understanding the distributions of potentially active TEs in the genome of 78 teleost fishes, representing major families and orders within the clade. Our analysis reveals coevolutionary arms race predicted by the Red Queen dynamics.

---

The jumping genes, also known as transposons or transposable elements (TE), jump from one locus to another and spread in the host genome. Because they can increase their copy number independent of host genome replication, they are also referred to as selfish genetic elements (Qiu et al. 2026; Meers et al. 2023). Increase in the TE copy number creates a metabolic burden for the host and their jump into regulatory and coding regions can substantially affect host survival and fitness. As a result, they are also considered as genomic parasites (Burt and Trivers 2006; Hayward and Gilbert 2022).

TE are categorised into two classes. Class I elements, also known as retrotransposons, follow a copy-and-paste mechanism for transposition, while Class II elements, also known as DNA transposons, follow cut-and-paste mechanism (Wicker et al. 2007; Bourque et al. 2018; Wells and Feschotte 2020). In Class I transposons the RNA transcript of a TE is converted into cDNA, which integrate itself in a different locus in the host genome increasing their copy number. While, in the case of Class II transposons, the RNA transcript is used for translating into a transposase that recognises the structural signatures of the original TE, cleaves it and integrates it into a different genomic locus (Shkumatov et al. 2022; Feschotte and Pritham 2007). Although the cut-and-paste mechanism apparently does not seem to increase the copy number, these TE often exploit the hemi-methylated state in the recently replicated genomic DNA and jump into yet to be replicated area of host genome near the replication fork. As a result, by taking the advantage of replication fork stalling the cut-and-paste TE can increase their copy number (Chen et al. 1987; Zhang et al. 2014; Ton-Hoang et al. 2010). Both Class I and Class II transposons can harbour non-autonomous elements, which failed to produce necessary enzymes for their transposition but they highjack enzymes produced by autonomous elements and jump (Hartl et al. 1992; Soundiramourtty and Mirouze 2025; Raiz et al. 2012; Wells and Feschotte 2020).

Increase in the TE copy number and their transposition in various loci in the genome creates a metabolic burden for the host and can sometimes leads to differential gene expression affecting the host survival and fitness (Burns 2022; Hayward and Gilbert 2022). As a result, hosts have evolved methods to regulate TE activity using a variety of mechanism including epigenetic silencing through DNA methylation and repressive histone modification, small RNA pathways including pi RNAs and RNAi, and protein mediated repression through KRAB zinc finger proteins and HUSH complex (Iwasaki et al 2025; Colonna and Fanti 2022; Davis et al., 2026; Lawlor and Ellison 2023).

Although host-TE co-existence is often discussed with respect to evolutionary arms race (Iwasaki et al. 2025; Qiu et al., 2026; Lawlor et al. 2023) and TE regarded as selfish genetic elements and genomic parasites (Qiu et al. 2026; Meers et al. 2023; Burt and Trivers 2006; Hayward and Gilbert 2022), there are recent arguments that refute selfish gene theory in favour of cooperation, co-option and mutualism (Cosby et al. 2019; Wang et al. 2022). Arguments against selfish gene theory regards TE as drivers of evolutionary change that help the host adapt to various environmental conditions (Fedoroff 2012; Schmidleithner, Stüve and Feuerer 2025). The mutualistic relationship between host and TE is often exemplified by contribution of TE to host survival in novel environments, change in host regulatory mechanisms to help various cellular activities, and domestication leading to evolution of novel traits (Ellison and Bachtrog 2013; Li et al. 2025; Ma et al. 2023; Modzelewski et al 2022). These arguments, although work in isolated examples, do not seem to be applicable to a wide variety of TE, which show a complete spectrum from extreme parasitism to mutualism (Kidwell and Lisch 2001). If the interaction between host and TE is mutualistic, we should expect their stable co-existence with minimum dynamics in the distribution pattern of TE in the host genome and host regulation against TE. If these traits continue to coevolve in the Red Queen dynamics in the evolutionary time scale, then it is an indication of co-evolutionary arms race with a continuous host-TE conflict.

Proposals of evolutionary arms race (Iwasaki et al. 2025; Qiu et al. 2026; Lawlor et al. 2023) and the red queen dynamics (McLaughlin and Malik 2017; Thomas-Bulle et al. 2018) in host-TE co-evolution have not been tested with a coherent hypothesis framework and testable predictions. Testing these predictions can have major impact on understanding host-TE conflict and distinguishing between dynamic arms race versus mutualism. We build a hypothesis testing framework (Fig. 1) to analyse the host-TE conflict in terms of the Red Queen dynamics. We build null and alternate hypothesis to understand how multilevel selection can lead to non-random distribution of TE with a preference for non-coding genic regions, variation in host response towards TE in evolutionary time and a test of Red Queen dynamics using a phylogenetic framework. To test this hypothesis, we studied distribution of potentially active structurally detected TE in 78 genomes representing major orders of the well supported Teleost clade (Fig. S1). Our analysis suggests that multilevel selection on TE distribution and dynamic nature of host regulation of TE follows Red Queen dynamics in the Teleost tree of life and likely explains the diversity of transposon distribution and regulation in all life forms in general.

**Fig. 1:**
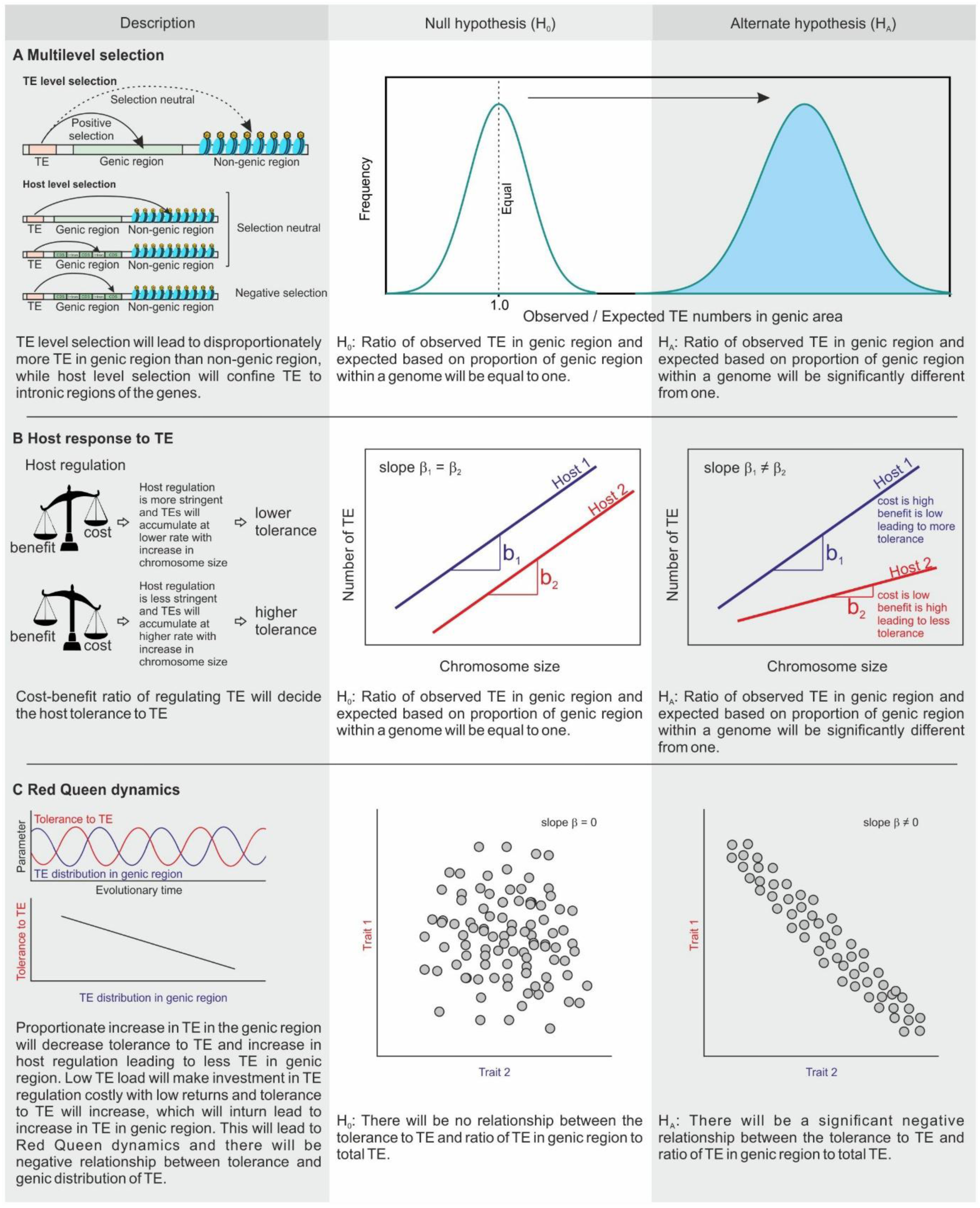
Hypothesis testing framework for the distribution of transposons (TE) in the genome, host regulation of TE, and Red Queen dynamics.

## Multilevel selection on TE distribution

Various genic regions within a genome like mRNA, tRNA, rRNA, small RNAs, lnc RNA, V and C gene regions, including pseudogenes (Pink et al. 2011; Wozniak et al., 2025) are often subject to transcriptional activation. TE that overlaps with these genes will have higher transcriptional activation than those present in non-genic regions of a genome. As a result, TE overlapping genic regions will have higher copy number and Darwinian fitness over transcriptionally inactivated TE in the non-genic regions. Therefore, selection acting at the level of TE will favour transposition of TE preferentially in the genic regions. However, if such transpositions compromise the fitness of the host by disrupting expression of genes essential for survival of the individual, host level selection will irradicate such transpositions. Multilevel selection, acting on the TE and on the host, will therefore affect the distribution of TE where they will be preferentially present in genic regions overlapping genes without affecting their function. This can be achieved by accumulation of TE in intronic regions of mRNA. Although alternating splicing and gene duplications will help the host survive TE transpositions in parts of essential genic regions, it is expected that most transpositions will accumulate in the intronic regions.

Above arguments can be tested by proposing null and alternating hypothesis (Fig. 1A). We will assume that fraction *f* of total genomic size in *bp* is occupied by various genic regions and 1–*f* by non-genic regions. If there are total *N* number of TE then under random distribution, we expect *Nf* number of TEs to be present in genic regions and *N*(1–*f*) in the non-genic regions. If the ratio of observed number of TE present in the genic region and the expected based on random distribution are equal, then across various genome we expect a normal distribution with mean of one. Alternatively, if the population mean is significantly different from one then the transposons are non-randomly distributed. If the ratio is more than one, it is an indication that transposons disproportionately overlap genic region than non-genic regions.

The ratio of observed and expected transposons in 78 teleost genomes followed a normal distribution (Shapiro-Wilk *W* = 0.98, *p* = 0.2599) with a mean 2.89 (S.E. 0.0529; range 1.56–3.92) (Fig. 2A). The mean was significantly different from one expected under random distribution (*t* = 35.77, *df* = 77, *p* < 0.0001) indicating that the distribution was non-random with higher preference for genic region. A similar argument can also be made with respect to the total size of TE in base pairs within genic and non-genic regions. Analysis of ratio of observed cumulative size of TE in the genic region versus expected under random distribution in genic region followed normal distribution (Shapiro-Wilk *W* = 0.9924, *p* = 0.9285) with population mean of 2.82 (S.E. 0.0528, range 1.71–3.89) (Fig. S2A). This mean was significantly different from expected under random distribution (*t* = 34.471, *df* = 77, *p* < 0.0001) indicating that the distribution was non-random with higher preference for genic region. Overlap of number of TE in various genic regions revealing their higher accumulation in introns (Fig. 2B) and the pattern was also consistent with the total size of the TE in *bp* (Fig. S2B).

**Fig. 2:**
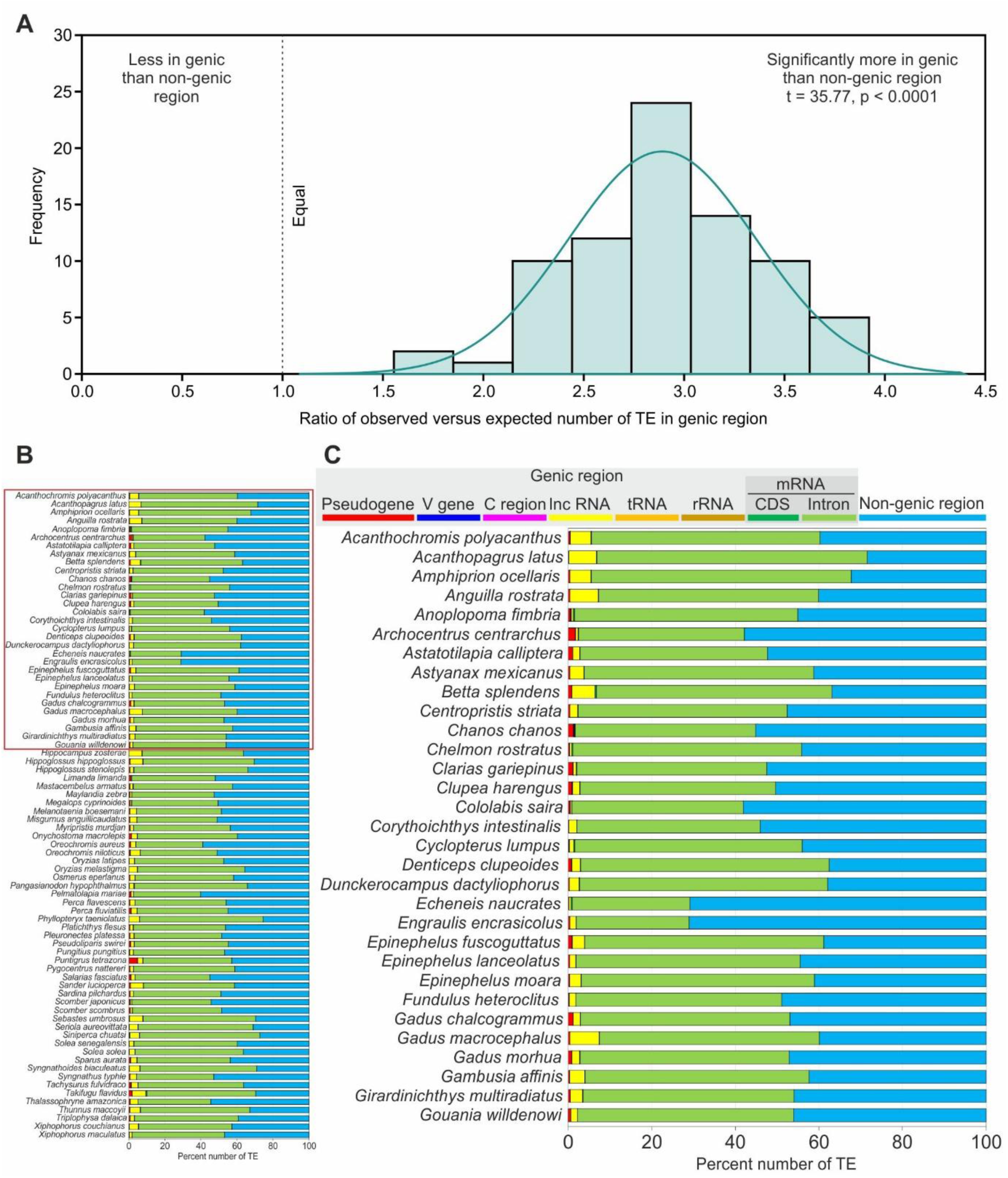
Distribution of TE in host genome. (A) Frequency of observed versus expected number of transposons in the genic region under random distribution. Significantly higher number of transposons in the genic region indicates preference of TE to genic region. (B) Distribution of TE in various genic region in 78 teleost fishes. (C) Selected view of (B) for higher resolution.

## Variability of host response against TE

Increase in the number of TE is an increasing function of genome size (Naville et al. 2019; Pedro et al. 2021). While this relationship provides how the TE numbers differ among species with different genome sizes, it does not provide information on the TE dynamics within a host. Since host genome is made of up chromosomes that vary in size, a relationship between chromosome size and the number of TE can provide information on how the host tolerate TE. If the relationship between the chromosome size and number of TE is linear, the slope of the line will provide information about how the TE accumulate with increase in the size of a chromosome within a host, providing information about tolerance of host to TE. The host response against TE, in terms of its tolerance, will be a function of cost to benefit ratio, of regulating TE. If different hosts have similar tolerance to TE, which is expected under mutualist relationship between host and TE, then the slopes of relationships between number of TE and chromosome size should not be significantly different among hosts, forming the null hypothesis. Alternatively, if the host and transposons are in conflict resulting in different cost benefit ratios the slope of the lines will differ significantly in some of the host speceis (Fig. 1B).

The relationships between number of TE and chromosome size in 78 Teleost species were significant and linear (Fig. 3A). For some of the organisms the slopes of the lines differed significantly. For example, the slopes of relationships in *Misgurunus anguillicaudatus* and *Cyclopterus lumpus* were significantly different (*t* = 30.71, *df* = 46, *p* < 0.0001). Similar arguments can be applied to cumulative size in *bp* of TE versus chromosome size in *bp* which also showed significant and linear relationships with differing slopes for different Teleost species (Fig. S3).

**Fig. 3:**
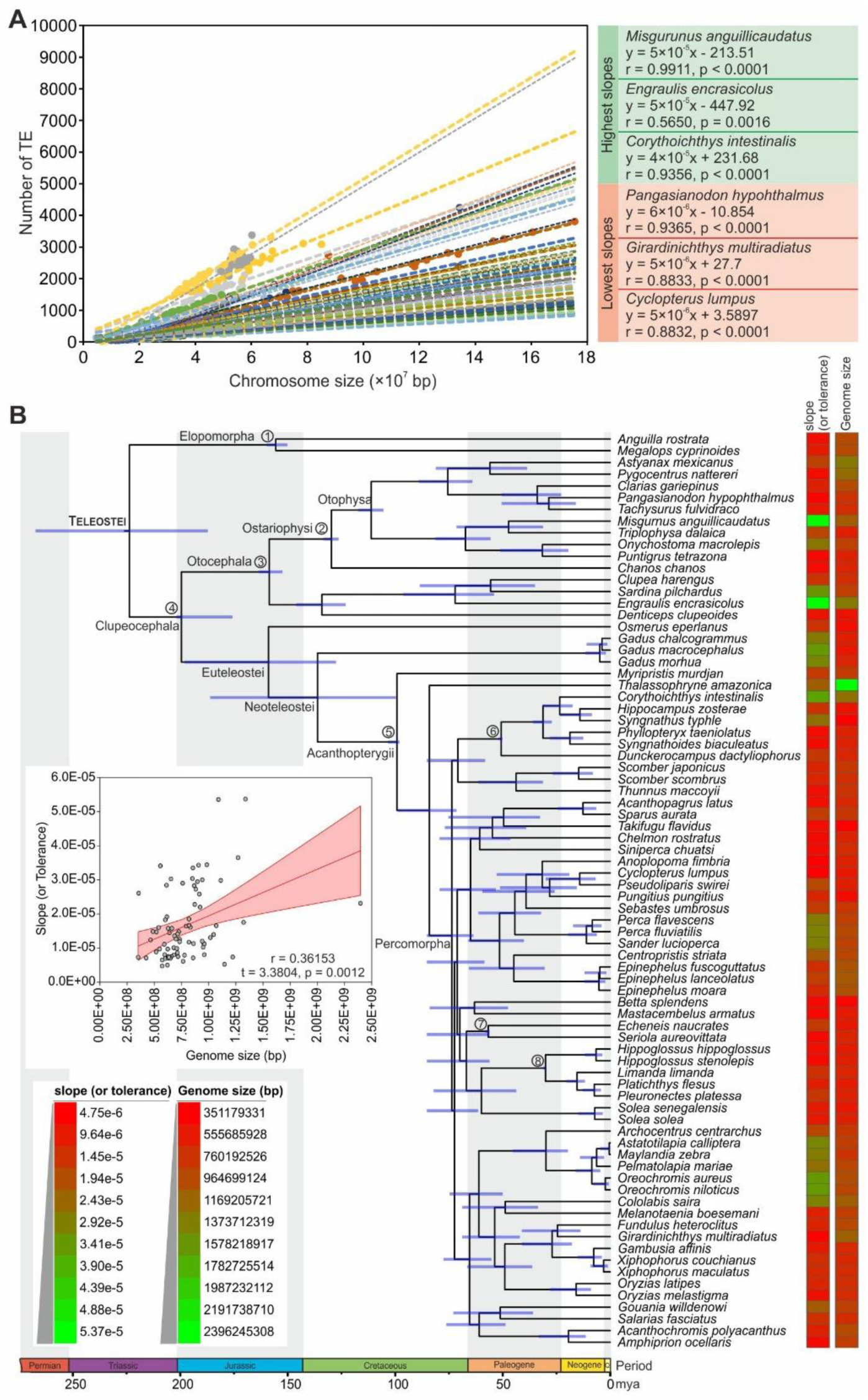
Host tolerance to TE numbers. (A) Relationship between the number of TE with chromosome size in 78 teleost species. Regression analysis for top three highest and bottom three lowest slopes sows marked variation in the slope (or tolerance) of the organisms to TE accumulation. (B) Fossil calibrated time tree depicting variation in the slope (or tolerance) in the Teleost fishes. There is a significant positive relationship between slope and genome size of an organism.

Accumulation of TE will likely increase the genome size and increase in the genome size will lead to higher cost of managing TE suggesting that higher tolerance to TE will be a function of genome size. We found a positive relationship between tolerance and genome size for both number of TE (Fig. 3B) and tolerance of cumulative size of TE (Fig. S3).

## Host-TE co evolution and the Red Queen dynamics

Increase of TE in the genic region is a fitness disadvantage for the host. As a result, increased investment of regulation of TE will fetch better benefits reducing the cost to benefit ratio. We therefore expect that, as the ratio of TE in genic region and total TE increases, the tolerance to TE should decrease. However, with less tolerance and high regulation by the host the TE numbers in the genic region will decrease. If the TE in the genic region decreases, investment in the high TE regulation by the host will not be cost effective increasing the cost to benefit ratio. As a result, host investment in regulation will decrease leading to increase in the tolerance. With increase in the tolerance to TE, the TE numbers in genic region will increase and the cycle will continue. This co-evolutionary arm race between host tolerance to TE and TE distribution in genic region, therefore, will follow the Red Queen dynamics (Fig. 1C).

Evolutionary arms race between the two traits, namely host tolerance to TE and TE distribution in genic region, can be tested against a null hypothesis that there are no relationships between the two traits, while the alternative hypothesis will predict a negative relationship under the Red Queen dynamics, where the decreases in one trait leads to increase in other and vice versa (Fig. 1C). Relationship between the tolerance to number of TE versus the ratio of number of TE in genic region out of total TE showed a significant (log-log plot, *r* = 0.5917, *t* = –6 .3992, *df* = 76, *p* < 0.0001) negative (slope *b* = –1.9819, S.E. = 0.3097, t = 6.3994, *df* = 76, *p* < 0.0001) relationship (Fig.4A). Above arguments can also be applied to the cumulative size of TE. Tolerance to cumulative size of TE and proportion of cumulative size of TE in genic region (Fig. S4A) showed a similar significant (log-log plot, r = –0.775, *t* = –67.181, *df* = 75, *p* < 0.0001) negative (slope *b* = –3.3476, S.E. = 0.04983, *t* = –67.18, *p* < 0.0001) relationship.

**Fig. 4:**
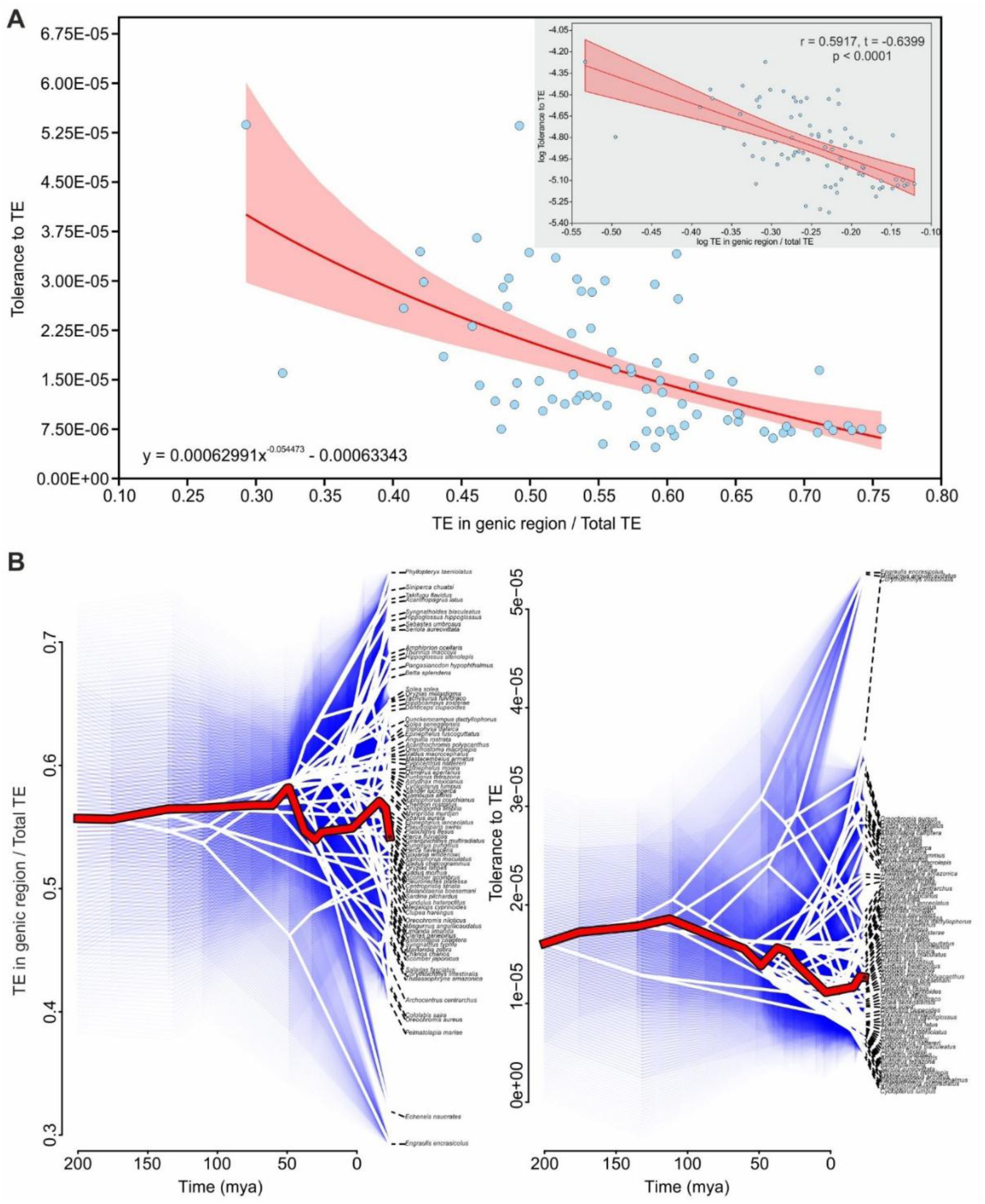
Host-TE arms race and the Red Queen dynamics. (A) Negative relationship between tolerance to TE and ratio of TE in genic and non-genic regions. (B) Phenogram of trait evolution with the red line indicating the evolution of trait in evolutionary time for *Xiphiphorous maculatus*, showing out of phase oscillations in the two traits.

To understand the probable change in the two traits in evolutionary time scale we reconstructed ancestral states in phylogeny to infer possible trait values on the branches (Fig. 5). There was significant phylogenetic signal for both tolerance to TE (Blomber’s K = 0.4024, *p* = 0.001) and the ratio of TE in genic region to total TE (Blomber’s K = 0.2609, *p* = 0.001) indicating that these traits have evolved along the phylogenetic tree. Because there was phylogenetic signal, we could reconstruct ancestral traits in phylogenies to understand trait evolution through time. Phenogram plotted by ancestral reconstruction showed out of phase oscillation in ancestral state reconstruction of the two trait values for different species. One example of the out of phase oscillation is provided for *Xiphiphorous maculatus* (Fig.4D).

**Fig. 5:**
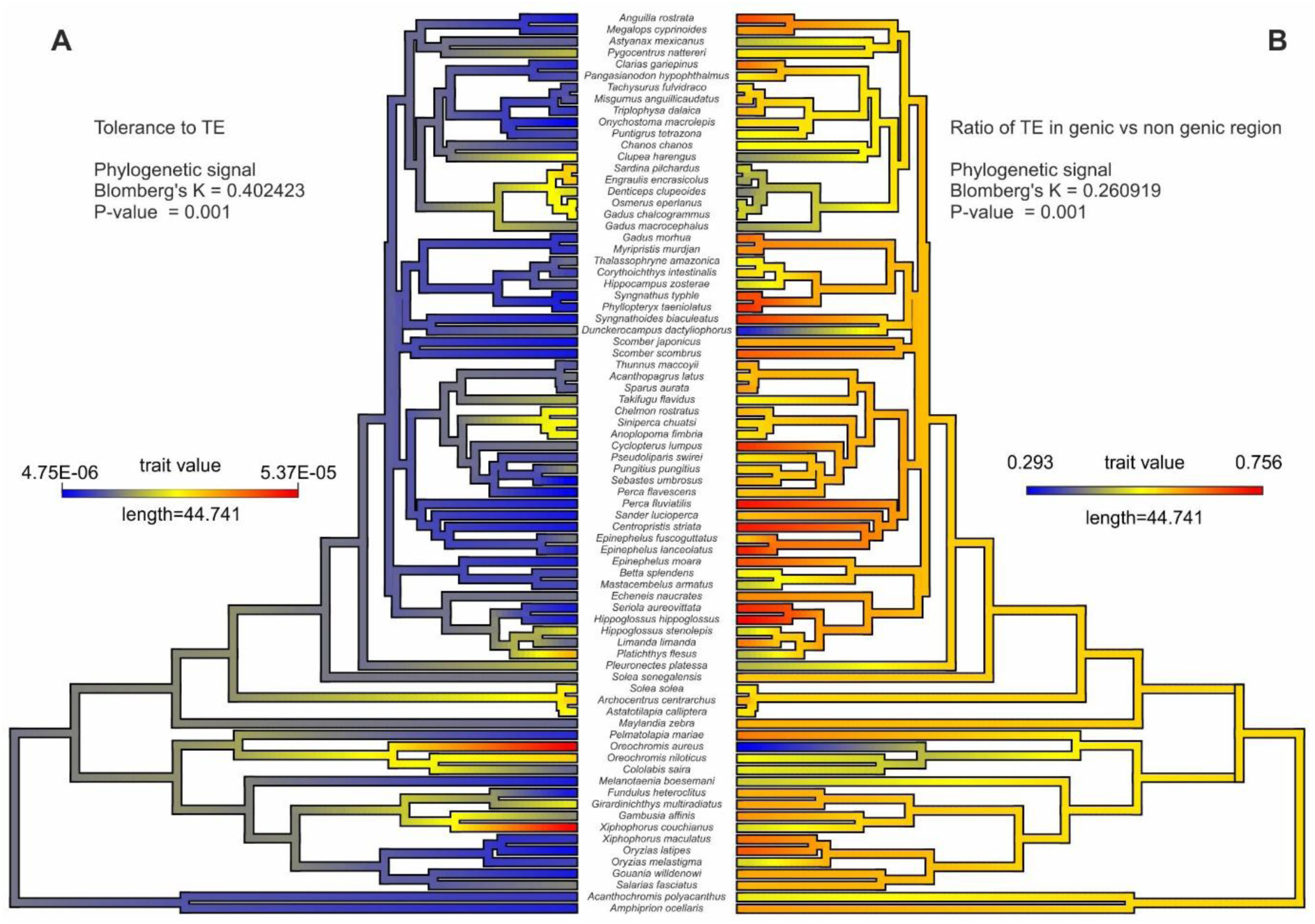
Ancestral trait reconstruction and phylogenetic signal analysis of host tolerance to TE and TE distribution in genic and non-genic regions. (A) Ancestral trait reconstruction for tolerance to TE. (B) Ancestral trait reconstruction for ratio of TE in genic versus non-genic regions. Both traits had significant phylogenetic signal on Teleost phylogeny.

## Discussion

The non-random distribution of transposons in the genome has already been suggested by earlier studies (Bourque et al. 2018; Wu et al. 2025; Paulat et al. 2022). However, most of the studies have argued that TE are more in non-genic regions and heterochromatin rather than in the genic areas (Quesneville 2020; Marsano et al. 2022; Hua-Van et al. 2005). On the contrary, we detected potentially active TE, that included structural features essential for their mobility, to be disproportionately more in the genic region than in the non-genic regions.

Studies that have argued higher proportion of TE in non-genic regions and heterochromatin included all detected TE using both structural as well as homology-based search methods. Homology based TE detection include TE lacking structural features that help it to transpose from one genomic locus to another. We expat that TE transposed in non-genic regions and heterochromatin are less likely to get transcribed. Further, these regions will be under neutral evolution and will accumulate mutations that can lead to loss of structural elements of a functional TE. Although these TE will retain homologous sequences, they will not contribute to copy number increase or TE fitness, which explains why heterochromatin are often referred as graveyards of TE (Bachmann 1996; Dimitri et al. 2009; Tamaru 2010; Šatović-Vukšić et al. 2025).

On the other hand, transposons that jumps into the genic region will have better fitness benefits due to the transcriptional activation (Capel et al.1993). These transposons will be under selection (Langmüller et al. 2023) and will retain their structural features. When we filtered out only structurally intact transposons, we showed that there are almost 3-fold more structurally active transposons in the genic regions than in non-genic regions clearly indicating a preference of TE to occupy genic regions. our results are in tuned with Zhang et al. (2020) who showed similar preference for genic regions by selected TE. Higher preference of TE for genic region can potentially affect host survival and fitness if they disrupt important functions. As a result, the selection acting on the host will result in a TE distribution pattern where TE are confined to non-codded regions such as introns (Wei et al. 2016; Paulat et al. 2022; Wu et al. 2025). Our observations are consistent with this suggestion in a wide range of teleost fishes. Although various previous studies have discussed the presence of TE in intronic regions, we argue that the disproportionate distribution of active TE in the genic region and their confinement to intronic regions is a result of multilevel selection acting at the level of TE and the host. Universality of these arguments in the 78 Teleost genomes, suggest that it should also be equally applicable to other taxa.

Regulation of TE by host shows wide variation across different taxonomic units (Pasquesi et al. 2020). Although methylation and small RNA mediated regulation is widespread in all organisms, considerable variation has been noted in how these mechanisms are executed for transposon control. Methylation status of repetitive elements differs among organisms (Zemach et al. 2010) and organisms such as plants, yeast and vertebrates have high methylation compared to protostomes, especially in insects (Canapa et al. 2015). Similarly, in the case of small RNA mediated response, although the principals of pathways are similar in various eukaryotic organisms, there are remarkable differences even in closely related species (Girard and Hannon 2008; Gebert and Rosenkranz 2015). Further, even within the same taxon, for example *Drosophila* (Canapa et al. 2015) and *Arabidopsis* (Jiang et al. 2024), it has been shown that the regulatory mechanism varies with transposon load, which has been argued to be a result of positive selection of regulatory mechanisms with increase in the threat of TE copy number increase. Counter mechanism from transposons to avoid host regulation also varies between different organisms. For example, in rice *Oryza sativa* a microRNA named as mir820 is present and carried by CACTA family of DNA transposons (Nosaka et al. 2012) can target OsDRM2 methyltransferase and reduce its activity leading to increase in the activity of transposons. In the case of *Arabidopsis*, siRNAs like tasiRNA, which is derived from Athila6 transposon, can target host defence mediated by UPB1b mRNA transcriptional repressor (McCue et al. 2013). In addition, VANDAL DNA transposable elements of *Arabidopsis* promote hypomethylation of own kind of transposons for aiding its transposition (Fu et al. 2013; Hosaka et al. 2017). The difference between host regulation strategies and TE counterattack strategies clearly shows there is a difference between regulation of transposon between different organisms (Cosby et al. 2019).

We have argued that this variation in the host response to the TE is an effect of cost-benefit trade-off. Increasing TE need not necessarily increase the regulation by the host. If the cost of regulating the TE is more than the benefit incur from regulating the TE host will learn to tolerate TE increase. On the other hand, if the benefit of controlling TE exceeds the cost of regulating TE host will regulate TE more stringently and lower tolerance to TE. This will lead to significant variation in the tolerance to TE among various hosts. We have provided a novel method to understand the tolerance by plotting number of TE versus the chromosome size for each organism. It is expected that number of TE will increase with chromosome size, but the rate of increase (explained by the slope of line) can infer about how much TEs are tolerated by organism with increase in the chromosome size. Larger is the slope, higher is the tolerance and vice versa. For the first time we show that within the teleost tree of life there is a considerable variation to the tolerance of TE. But we expect these trends will be universal for other taxa.

The Red Queen dynamics between the host and the transposon has been argued in a variety of scenarios (McLaughlin and Malik 2017; Thomas-Bulle et al. 2018; Kosuge et al. 2024). Change in the TE assemblage, due to horizontal transfer of TE in *Drosophila*, lead to adaptive evolution in the piRNA machinery to curb the TE transposition leading to a co-evolutionary arms race predicted by Red Queen dynamics (Blumenstiel et al. 2016; Parhad et al. 2017; Luo et al. 2020). Similarly, the expansion of the Krüppel-associated box-containing zinc finger proteins (KRAB-ZFPs), which epigenetically suppress TE to counteract disorderly TE transpositions, has been attributed to the evolution of TE families (Kosuge and Hamada 2024).

Our argument of Red Queen works at the phylogenetic level. We argue that if there are less TE in the genic region, compared to the non-genic region, the host fitness is not compromised substantially and the cost of investment in regulating the TE for host will surpass the benefit incurred. Under such condition host will have low regulation leading to higher tolerance to TE increase. However, under these conditions TE will start accumulating in genic region, where they will have selective advantage for increasing their copy number leading to increase in the TE load. This will lead to accumulation of TE disproportionately more in the genic versus non-genic region. This can substantially affect the host fitness and controlling TE will be more beneficial compared to the cost of regulation, leading to decrease in the tolerance and most stringent regulation of TE by the host. This can substantially reduce the TE number especially in the genic region. Reduce number of TEs makes investment in TE regulation futile, which will increase the tolerance host to TE and the cycle will continue. These dynamics will lead to a coevolutionary arms race between host regulation and TE distribution. The implied negative relationship between the tolerance and TE in genic versus non-genic region fits perfectly with the data obtained from teleost fishes. Further, by reconstruing the ancestral state in phylogenies, we showed that these two traits oscillate in evolutionary time scale. Because our arguments are in general, we expect similar trends to occur for other taxa as well.

We considered only structurally detected TE for analysis as their structural integrity vouch for their ability to transpose, affecting their fitness through increase in the copy number. Transposons that were detected only based on homology lacked the structural features essential for transposition and therefore they were ignored from analysis. Some of the DNA transposons, like Polinton, Merlin, PiggyBac, Sola1, Soka2 and P elements, which were only detected based on homology, could not be considered in the final analysis. Further, two retrotransposons, Line and Penelope which lack well defined structural features like LTR (Craig et al. 2021; Ghanim et al. 2025), were also detected only based on homology and were excluded from analysis. However, exclusion of these TE is less likely to affect our results qualitatively as they collectively contributed to jut about 3.5% of total number TE that were detected. It is also essential to note that, along with exact boundaries of gene several flanking regions of genes also come under open chromatin structures hence can be transcriptionally active. Studies have shown that jumping Transposons can get inserted into these genomic locations (Wei et al. 2016; Wu et al. 2025). These locations will come under intergenic regions. In our study, we did not consider much flanking regions of the genes. Although TE overlapping starting and end of genes were considered, TE present in the flanking regions were not considered as part of overlapping the genic region. This have led to grouping of TE in these regions under non-genic region instead of genic region. However, we do not expect it to affect statistical results that we have shown as is only hints to an underestimation.

The continual evolutionary arms race in host regulation and TE distribution suggests that viewing TE- host coexistence as mutualism may not be appropriate. Although in selected examples TE have been shown to enhance host fitness either by performing host beneficial traits or through domestication. Such cases are relatively rare and may not portray large scale distribution of transposons among hosts.

## Acknowledgements

We are thankful to Sanjeev Galande for continuous support and encouragement and to Milind Watve for helpful discussion. Ashish Gupta and Ashitosh Singh provided valuable comments during the work progress. We thank Shiv Nadar Institution of Eminence, Delhi-NCR, for providing infrastructural facilities. MP is thankful to the Shiv Nadar Institution of Eminence, Delhi-NCR, for doctoral fellowship.

## Author contribution

Conceptualization: MP, ND

Methodology: MP, ND

Software: MP, SP

Validation: MP, ND

Formal analysis: MP, ND

Investigation: MP, SP

Resources: ND

Data Curation: MP, ND

Writing - Original Draft: MP, ND

Writing - Review & Editing: ND

Visualization: MP, ND

Supervision: ND

## Reference

Bachmann, L. (1996) Evolutionary changes of the structure of mobile genetic elements in Drosophila. In: Tomiuk, J., Wöhrmann, K., Sentker, A. (eds) Transgenic Organisms. Advances in Life Sciences. Birkhäuser Basel. 10.1007/978-3-0348-9177-6_4

Blumenstiel, J.P., Erwin, A.A. and Hemmer, L.W. (2016) What drives positive selection in the *Drosophila* piRNA machinery? The genomic autoimmunity hypothesis. The Yale Journal of Biology and Medicine, 89(4), 499–512.

Bourque, G., Burns, K.H., Gehring, M., Gorbunova, V., Seluanov, A., Hammell, M., Imbeault, M., Izsvák, Z., Levin, H.L., Macfarlan, T.S. and Mager, D.L. (2018) Ten things you should know about transposable elements. Genome Biology, 19, 199. 10.1186/s13059-018-1577-z

Burns, K. H. (2022) Repetitive DNA in disease. Science, 376, 353–354. 10.1126/science.abl739

Burt, A. and Trivers, R. (2006) *Genes in conflict: the biology of selfish genetic elements*. Harvard University Press.

Canapa, A., Barucca, M., Biscotti, M.A., Forconi, M. and Olmo, E. (2015) Transposons, genome size, and evolutionary insights in animals. Cytogenetic and Genome Research, 147, 217–239. 10.1159/000444429

Capel, J., Montero, L.M., Martínez-Zapater, J.M. and Salinas, J. (1993) Non-random distribution of transposable elements in the nuclear genome of plants. Nucleic Acids Research, 21, 2369–2373. 10.1093/nar/21.10.2369

Chen, J., Greenblatt, I.M. and Dellaporta, S.L. (1987) Transposition of Ac from the P locus of maize into unreplicated chromosomal sites. Genetics, 117, 109–116. 10.1093/genetics/117.1.109

Colonna Romano, N. and Fanti, L. (2022) Transposable elements: major players in shaping genomic and evolutionary patterns. Cells, 11, 1048. 10.3390/cells11061048

Cosby, R.L., Chang, N.C. and Feschotte, C. (2019) Host–transposon interactions: conflict, cooperation, and cooption. Genes and Development, 33, 1098–1116. 10.1101/gad.327312.119

Craig, R.J., Yushenova, I.A., Rodriguez, F. and Arkhipova, I.R. (2021) An ancient clade of Penelope-like retroelements with permuted domains is present in the green lineage and protists, and dominates many invertebrate genomes. Molecular Biology and Evolution, 38, 5005–5020. 10.1093/molbev/msab225

Davis, J., Voicu, D., Chitnavis, U., Jaksina, J. and Imbeault, M. (2026) The role of KRAB zinc-finger proteins in expanding the domestication potential of transposable elements. Nature Genetics, 58, 492–502. 10.1038/s41588-025-02498-3

Dimitri, P., Caizzi, R., Giordano, E., Carmela Accardo, M., Lattanzi, G. and Biamonti, G., (2009) Constitutive heterochromatin: a surprising variety of expressed sequences. Chromosoma, 118, 419–435. 10.1007/s00412-009-0211-y

Ellison, C.E. and Bachtrog, D. (2013) Dosage compensation via transposable element mediated rewiring of a regulatory network. Science, 342, 846–850. 10.1126/science.1239552

Fedoroff, N.V. (2012) Transposable elements, epigenetics, and genome evolution. Science, 338, 758–767. 10.1126/science.338.6108.758

Feschotte, C. and Pritham, E.J. (2007) DNA transposons and the evolution of eukaryotic genomes. Annual Review of Genetics, 41, 331–368. 10.1146/annurev.genet.40.110405.090448

Fu, Y., Kawabe, A., Etcheverry, M., Ito, T., Toyoda, A., Fujiyama, A., Colot, V., Tarutani, Y. and Kakutani, T. (2013) Mobilization of a plant transposon by expression of the transposon-encoded anti-silencing factor. EMBO Journal, 32, 2407–2417. 10.1038/emboj.2013.169

Gebert, D. and Rosenkranz, D. (2015) RNA-based regulation of transposon expression. Wiley Interdisciplinary Reviews: RNA, 6, 687–708. 10.1002/wrna.1310

Ghanim, G.E., Hu, H., Boulanger, J. and Nguyen, T.H.D. (2025) Structural mechanism of LINE-1 target-primed reverse transcription. Science, 388, eads8412. 10.1126/science.ads8412

Girard, A. and Hannon, G.J. (2008) Conserved themes in small-RNA-mediated transposon control. Trends in Cell Biology, 18, 136–148. 10.1016/j.tcb.2008.01.004

Hartl, D.L., Lozovskaya, E.R. and Lawrence, J.G. (1992) Nonautonomous transposable elements in prokaryotes and eukaryotes. Genetica, 86, 47–53.

Hayward, A. and Gilbert, C. (2022) Transposable elements. Current Biology, 32, R904–R909. 10.1016/j.cub.2022.07.044

Hosaka, A., Saito, R., Takashima, K., Sasaki, T., Fu, Y., Kawabe, A., Ito, T., Toyoda, A., Fujiyama, A., Tarutani, Y. and Kakutani, T. (2017) Evolution of sequence-specific anti-silencing systems in *Arabidopsis*. Nature Communication, 8, 2161. 10.1002/wrna.70022

Hua-Van, A., Le Rouzic, A., Maisonhaute, C. and Capy, P. (2005) Abundance, distribution and dynamics of retrotransposable elements and transposons: similarities and differences. Cytogenetic and Genome Research, 110, 426–440. 10.1159/000084975

Iwasaki, Y.W., Shoji, K., Nakagwa, S., Miyoshi, T. and Tomari, Y. (2025) Transposon–host arms race: a saga of genome evolution. Trends in Genetics, 41, 369–389. 10.1016/j.tig.2025.01.009

Jiang, J., Xu, Y.C., Zhang, Z.Q., Chen, J.F., Niu, X.M., Hou, X.H., Li, X.T., Wang, L., Zhang, Y.E., Ge, S. and Guo, Y.L. (2024) Forces driving transposable element load variation during Arabidopsis range expansion. The Plant Cell, 36, 840–862. 10.1093/plcell/koad296

Kidwell, M.G. and Lisch, D.R. (2001) Perspective: transposable elements, parasitic DNA, and genome evolution. Evolution, 55, 1–24. 10.1111/j.0014-3820.2001.tb01268.x

Kosuge, M., Ito, J. and Hamada, M. (2024) Landscape of evolutionary arms races between transposable elements and KRAB-ZFP family. Scientific Reports, 14, 23358. 10.1038/s41598-024-73752-7

Langmüller, A.M., Nolte, V., Dolezal, M. and Schlötterer, C. (2023) The genomic distribution of transposable elements is driven by spatially variable purifying selection. Nucleic Acids Research, 51, 9203–9213. 10.1093/nar/gkad635

Lawlor, M.A. and Ellison, C.E., 2023. Evolutionary dynamics between transposable elements and their host genomes: mechanisms of suppression and escape. Current Opinion in Genetics & Development, 82, 102092. 10.1016/j.gde.2023.102092

Li, T.D., Toohill, K. and Modzelewski, A.J. (2025) From junk DNA to genomic treasure: impacts of transposable element DNA, RNA, and protein in mammalian development and disease. Wiley Interdisciplinary Reviews: RNA, 16, e70022. 10.1002/wrna.70022

Luo, S., Zhang, H., Duan, Y., Yao, X., Clark, A.G. and Lu, J. (2020) The evolutionary arms race between transposable elements and piRNAs in *Drosophila* melanogaster. BMC Evolutionary Biology, 20, 14. 10.1186/s12862-020-1580-3

Ma, H., Wang, M., Zhang, Y.E. and Tan, S. (2023) The power of “controllers”: Transposon-mediated duplicated genes evolve towards neofunctionalization. Journal of Genetics and Genomics, 50, 462–472. 10.1016/j.jgg.2023.04.003

Marsano, R.M. and Dimitri, P. (2022) Constitutive heterochromatin in eukaryotic genomes: A mine of transposable elements. Cells, 11, 761. 10.3390/cells11050761

McCue, A.D., Nuthikattu, S. and Slotkin R K. (2013) Genome-wide identification of genes regulated in trans by transposable element small interfering RNAs. RNA Biology 10: 1379–1395. 10.4161/rna.25555

McLaughlin Jr, R.N. and Malik, H.S. (2017) Genetic conflicts: the usual suspects and beyond. Journal of Experimental Biology, 220, 6–17. 10.1242/jeb.148148

Meers, C., Le, H.C., Pesari, S.R., Hoffmann, F.T., Walker, M.W., Gezelle, J., Tang, S. and Sternberg, S.H. (2023) Transposon-encoded nucleases use guide RNAs to promote their selfish spread. Nature, 622, 863–871. 10.1038/s41586-023-06597-1

Modzelewski, A.J., Gan Chong, J., Wang, T. and He, L. (2022) Mammalian genome innovation through transposon domestication. Nature Cell Biology, 24, 1332–1340. 10.1038/s41556-022-00970-4

Naville, M., Henriet, S., Warren, I., Sumic, S., Reeve, M., Volff, J.N. and Chourrout, D. (2019) Massive changes of genome size driven by expansions of non-autonomous transposable elements. Current Biology, 29, 1161–1168. 10.1016/j.cub.2019.01.080

Nosaka, M., Itoh, J.-I., Nagato, Y., Ono, A., Ishiwata A. and Sato Y. (2012. Role of transposon-derived small RNAs in the interplay between genomes and parasitic DNA in rice. PLoS Genetics 8: e1002953. 10.1371/journal.pgen.1002953

Parhad, S.S., Tu, S., Weng, Z. and Theurkauf, W.E. (2017) Adaptive evolution leads to cross-species incompatibility in the piRNA transposon silencing machinery. Developmental Cell, 43, 60–70. 10.1016/j.devcel.2017.08.012

Pasquesi, G.I., Perry, B.W., Vandewege, M.W., Ruggiero, R.P., Schield, D.R. and Castoe, T.A. (2020) Vertebrate lineages exhibit diverse patterns of transposable element regulation and expression across tissues. Genome Biology and Evolution, 12, 506–521. 10.1093/gbe/evaa068

Paulat, N.S., McGuire, E., Subramanian, K., Osmanski, A.B., Moreno-Santillán, D.D., Ray, D.A. and Xing, J. (2022) Transposable elements in bats show differential accumulation patterns determined by class and functionality. Life, 12, 1190. 10.3390/life12081190

Pedro, D.L.F., Amorim, T.S., Varani, A., Guyot, R., Domingues, D.S. and Paschoal, A.R., 2021. An atlas of plant transposable elements. F1000Research, 10, 1194. 10.12688/f1000research.74524.1

Pink, R.C., Wicks, K., Caley, D.P., Punch, E.K., Jacobs, L. and Francisco Carter, D.R. (2011) Pseudogenes: pseudo-functional or key regulators in health and disease? RNA, 17, 792–798. 10.1261/rna.2658311

Qiu, B., Elsner, D. and Korb, J. (2026) Arms races between selfish genetic elements and their host defence in termites. Nature Communications, 17, 1702. 10.1038/s41467-026-69550-6

Quesneville, H. (2020) Twenty years of transposable element analysis in the *Arabidopsis* thaliana genome. Mobile DNA, 11, 28. 10.1186/s13100-020-00223-x

Raiz, J., Damert, A., Chira, S., Held, U., Klawitter, S., Hamdorf, M., Löwer, J., Strätling, W.H., Löwer, R. and Schumann, G.G. (2012) The non-autonomous retrotransposon SVA is trans-mobilized by the human LINE-1 protein machinery. Nucleic Acids Research, 40, 1666–1683. 10.1093/nar/gkr863

Šatović-Vukšić, E., Majcen, P. and Plohl, M. (2025) Satellite DNAs rising from the transposon graveyards. DNA Research, 32, dsaf026. 10.1093/dnares/dsaf026

Schmidleithner, L., Stüve, P. and Feuerer, M. (2025) Transposable elements as instructors of the immune system. Nature Reviews Immunology, 25, 696–706. 10.1038/s41577-025-01172-3

Shkumatov, A.V., Aryanpour, N., Oger, C.A., Goossens, G., Hallet, B.F. and Efremov, R.G., (2022) Structural insight into Tn3 family transposition mechanism. Nature Communications, 13, 6155. 10.1038/s41467-022-33871-z

Soundiramourtty, A. and Mirouze, M. (2025) Plant MITEs: miniature transposable elements with major impacts. Mobile DNA, 16, 43. 10.1186/s13100-025-00375-8

Tamaru, H. (2010) Confining euchromatin/heterochromatin territory: jumonji crosses the line. Genes & Development, 24, 1465–1478. http://www.genesdev.org/cgi/doi/10.1101/gad.1941010.

Thomas-Bulle, C., Piednoël, M., Donnart, T., Filée, J., Jollivet, D. and Bonnivard, É. (2018) Mollusc genomes reveal variability in patterns of LTR-retrotransposons dynamics. BMC Genomics 19, 821. 10.1186/s12864-018-5200-1

Ton-Hoang, B., Pasternak, C., Siguier, P., Guynet, C., Hickman, A.B., Dyda, F., Sommer, S. and Chandler, M. (2010) Single-stranded DNA transposition is coupled to host replication. Cell, 142, 398–408. 10.1016/j.cell.2010.06.034

Wang, S., Koide, Y. and Kishima, Y. (2022) How to establish a mutually beneficial relationship between a transposon and its host: lessons from Tam3 in Antirrhinum. Genes & Genetic Systems, 97, 177–184. 10.1266/ggs.22-00063

Wei, B., Liu, H., Liu, X., Xiao, Q., Wang, Y., Zhang, J., Hu, Y., Liu, Y., Yu, G. and Huang, Y. (2016) Genome-wide characterization of non-reference transposons in crops suggests non-random insertion. BMC Genomics, 17, 536. 10.1186/s12864-016-2847-3

Wells, J.N. and Feschotte, C. (2020) A field guide to eukaryotic transposable elements. Annual Review of Genetics, 54, 539–561. 10.1146/annurev-genet-040620-022145

Wicker, T., Sabot, F., Hua-Van, A., Bennetzen, J.L., Capy, P., Chalhoub, B., Flavell, A., Leroy, P., Morgante, M., Panaud, O. and Paux, E. (2007) A unified classification system for eukaryotic transposable elements. Nature Reviews Genetics, 8, 973–982. 10.1038/nrg2165

Wozniak, J., Stachowiak, M., Switonski, M. and Nowacka-Woszuk, J. (2025) Transcript patterns of bovine CYP21A2 and its pseudogene in adrenal and ovarian tissues. Genes, 16, 1374. 10.3390/genes16111374

Wu, W., Zeng, Y., Huang, Z., Peng, H., Sun, Z. and Xu, B. (2025) Transposable element landscape in the monotypic species Barthea barthei (Hance) Krass (Melastomataceae) and its role in ecological adaptation. Biomolecules, 15, 346. 10.3390/biom15030346

Zemach, A., McDaniel, I.E., Silva, P. and Zilberman, D. (2010) Genome-wide evolutionary analysis of eukaryotic DNA methylation. Science, 328, 916–919. 10.1126/science.1186366

Zhang, J., Zuo, T., Wang, D. and Peterson, T. (2014) Transposition-mediated DNA re-replication in maize. Elife, 3, e03724. 10.7554/eLife.03724

Zhang, X., Zhao, M., McCarty, D.R. and Lisch, D. (2020) Transposable elements employ distinct integration strategies with respect to transcriptional landscapes in eukaryotic genomes. Nucleic Acids Research, 48, 6685–6698. 10.1093/nar/gkaa370

