## Supplementary material for "Evolutionary genomics of host-transposon conflict, multilevel selection, and Red Queen dynamics": Fig. S1

**The PDF file includes:**

Materials and Methods

Figs. S1 to S3

### Materials and Methods

#### *Data acquisition*

To test this hypothesis, we needed to count the TE number and total occupancy in various Genomes. We selected entire teleost fish clade to perform this analysis, and the method use to collect the data is explained below. The National Centre for Biotechnology Information (NCBI) (https://www.ncbi.nlm.nih.gov/) was scanned for all available assembled and annotated Teleost fish genomes. A total of 78 genomes of representative species for all Teleost fish having chromosome level assembly and complete annotation were chosen for the analysis. The GFF (having cytological map position for different genic feature) along with Fasta sequence of these organisms were downloaded from NCBI. The genome size for each of these species were recalculated as the sum of all chromosome size.

#### *Annotation of TEs*

The Extensive De Novo TE Annotator pipeline or EDTA (Ou et al 2019) was used to scan for the presence of Transposable elements (TE) in these 78 Genomes. The EDTA pipeline have eight gold standard Tools for TE detection. For LTR (Long terminal repeats) retrotransposons detection tools like LTRharvest (Ellinghaus *et al.* 2008), LTR_FINDER_parallel (Xu and Wang 2007) and LTR_retriever (Ou and Jiang 2018) are used whereas Generic Repeat Finder (Shi and Liang 2019), TIR-Learner (Su, Gu and Peterson 2019.) and HelitronScanner (Xiong et al. 2014.) are used for TIR transposons and helitron detection respectively. Along with these structure-based methods RepeatModeler (Smit, Hubley and Green 2015) is used to predict non-LTR retrotransposons SINEs and LINEs. EDTA uses RepeatMasker (Smit, Hubley and Green 2015) to annotate fragmented TEs and masking of repeat regions. EDTA can detect intact and full length Transposoons which have completed structural signatures and significant protein domains. EDTA can also detect those elements which are fractured or fragmented. These elements either have partial structural signatures or do not have and structural signature but shows sequence similarity with known TE proteins.

We have used EDTA for non-model organisms and with parameters –species others, --species run the entire pipeline(all), --sensitive (RepeatModeler) –a --evaluate. EDTA can detect intact and full-length Transposons which have completed structural signatures and significant protein domains.

For example, DNA (TIR) type of TE has two terminal inverted repeats (TIRs) at both of its end embedding a single gene to produce Transposase enzyme (Tpase). They generate Terminal site duplications (TSD) during their insertions. DNA(Helitron) type of TE encodes RPA (replication protein A) and to transpose they follow a rolling circle replication (Kapitonov and Jurka 2007). They contain a ZN (zinc) finger domain followed by replication protein A (REP) like domain and helicase like domain (HEL). TC sequence followed by 22nt AT rich region in the 5' terminal and GC rich hairpin like structure in the 3' end are marked as structural signatures Helitrons (Thomas and Pritham 2015).

Another type DNA TE is known as POLINTON /Mavericks (largest type of transposons). They are characterized with perfect terminal inverted repeats and can code for up to 10 genes. They can encode for DNA polymerase by themselves. During insertion they induces terminal site duplications (Kapitonov, and Jurka, 2006). Structural signatures of these elements can be characterized by long terminal inverted repeats containing DNA polymerase like enzymes in them.

In case of RNA type of TE EDTA can efficiently detect LTRs (Long Terminal Repeats). These elements have very long terminal repeats enclosing several enzymes for copying itself. there is a primer binding site (PBS) immediately after the LTR. The internal region after the LTR is responsible to code mostly two genes i.e., gag and pol. A single protein known as capsid-like protein is encoded by the gag gene whereas the pol is known to encode a poly protein that shows reverse transcriptase (RT) activity, protease activity RNase H, and integrase activity responsible for successful transposition reaction (Xu et al. 2010).

Another type of RNA TE is described as NON LTRs. These are the retro transposons which do not possess the long terminal inverted repeats (LTR). They are also known as polyA retrotransposons, NON LTRs or target-primed (TP) retrotransposons). Instead of having LTRs on the ends it shows property like a genome integrated mRNA (Han, 2010). These elements can be long up to several kb. They do not have LTRs, but they have a poly A tail. It is consisting of two ORFs. The ORF 1 codes for gag and the ORF codes for Endo nuclease (EN), reverse transcriptase (RT). These ORFs are embedded inside two UTRs i.e., the 5'UTR and 3'UTR. Followed by 3'UTR of the structure a poly A sequence is there (Grechishnikova and Poptsova 2016).

Along with these full length TEs non- autonomous elements showing jumping signatures can also be detected by EDTA with structural method of TE detection.

EDTA can also detect those elements which are fractured or fragmented. These elements either have partial structural signatures or do not have and structural signature but shows sequence similarity with known TE proteins (Ou *et al*. 2019). We have considered only those TEs which have intact structural signatures (detected by structural methods) and have potential to jump (potentially active TEs).

Several Retrotransposons specifically non-LTR retrotransposons and Pointon were either not detected or rarely detected by Structural methods-based TE identification, so we did not consider those elements for further analysis. For this analysis we have considered DNA/DTA, MITE/DTA, DNA/DTC, MITE/DTC, DNA/DTH, MITE/DTH, DNA/DTM, MITE/DTM, DNA/DTT, MITE/DTT, DNA/Helitron, LTR/Copia, LTR/Gypsy, LTR/unknown as they have shown significant jumping signature. We have used usegalaxy.eu along with

EDTA for the whole Genome TE annotation. A total of 28 orders, 78 genera, 78 species from Teleostei were annotated and analysed for transposable element distribution study.

***Distribution study***

We choose structurally identified TEs for further analysis. The annotation file generated out of the EDTA contain positional information of detected TEs. We have compared these data with the genic and intergenic positions of the whole genome to study the distribution of these elements around the genes. We have used several self-designed simple python scripts to extract co-ordinates of those TE which have overlap with the gene feature described in NCBI whole genome annotation (GFF3) file. The Genome annotation file of NCBI have cytologic positions for each gene in the Genome. In the files the Genes can be mRNA, rRNA, tRNA, LncRNA, snRNA, snoRNA, Pseudogenes, V gene region and C gene region. We have used a logic where the code searches for TEs, those have either starting position (TE start) greater than the genomic features starting position (Feature start) and the ending positions (TE end) less than the genomic features end position (Feature end) or TE start less than Feature start and TE end is less than Feature end or TE start is greater than Feature start and TE end is greater than Feature end. For mRNA we have searched TE overlap on both exons and introns with same logic described above. If we found any TE overlapping exon partially and overlapping intron partially those will be considered as Exon overlapping TEs. Finally, the extracted information's were analysed by several statistical analysis. Similar workflow was also used to see how fragmented and not intact transposons (detected by homology-based method) was distributed in the mRNA and Pseudogenes.

***Phylogenetic tree construction***

To trace the phylogenetic relationship among the 78 teleost species we used amino acid sequences of 51 orthologous genes. We used Benchmarking Universal Single-Copy Orthologs, BUSCO (Waterhouse et al. 2018) for identification of 51 single copy orthologous genes that were common among 78 species. As they were in single copy and common across all these 78 genomes, we have collected the amino acid sequences of these proteins (generated during BUSCO analysis) to build the phylogenetic tree. The amino acid sequences for each gene were separated out for every organism and same genes for same organism were kept in single file. molecular evolutionary genetics analysis (MEGA) and MUSCLE was used to do multiple sequence alignments (MSA) (Tamura et al 2007, Edgar 2004). The MSA were further edited with using Gblocks (Castresana 2000). Followed by editing IQ-TREE was used to create the phylogeny based on these 51 genes (Nguyen et al 2015). For time tree calibration points we have used the data available in Hughes et al 2018, Ghezelayagh et al 2022 and Betancur-R et al. 2013. the constraints we have used are as follows.

1. Elopomorpha. MRCA: Megalops, Anguilla. Hard minimum age: 149 Ma. 95% soft maximum age: 250 Ma. Calibration source (Broughton et al. 2013).
2. Ostariophysi. MRCA: Chanos, Danio. Hard minimum age: 126.3 Ma. 95% soft maximum age: 158.3 Ma. Calibration source (Benton et al. 2015).
3. Otomorpha. MRCA: Amblygaster, Danio. Hard minimum age: 150.94 Ma. 95% soft maximum age: 228.4 Ma. Calibration source (Benton et al. 2015).
4. Clupeocephala. MRCA: Danio, Lepidogalaxias. Hard minimum age: 150.94 Ma. 95% soft maximum age: 235 Ma. Calibration source (Benton et al. 2015 45).
5. Holocentriformes+Percomorphaceae. MRCA: Sargocentron, Aulostomus. Hard minimum age: 98.0 Ma. 95% soft upper bound: 128.8 Ma. Calibration source (Harrington et al. 2016 50).

6. Syngnathidae. MRCA: Syngnathus, Syngnathoides. Hard minimum age: 50 Ma. 95% soft maximum age: 57.5 Ma. Calibration source (Near et al. 2012, modified by Betancur-R. et al. 2013).
7. Stem lineage Pleuronectidae, dating the MRCA of Pleuronichthys cornutus and Paralichthys albigutta, which subtends the MRCA of the clade containing Pleuronectidae and Paralichthyidae. First occurrence: †Oligopleuronectes germanicus from the Frauenweiler fossil site, Germany109. Resolution in phylogenetic analyses: None. Character states: Suggested as a member of Pleuronectidae based on †Oligopleuronectes being right-eyed and having a lateral process on the eye-side frontal109,110. Stratigraphy: See discussion in Harrington et al.1. Absolute age estimate: 29.62 Ma1. Prior setting: lognormal prior, minimum age 29.62 Ma, mean=1.0, S.D.= 0.45, upper 95% CI: 35.3 Ma1.
8. Carangiformes. MRCA: Caranx, Echeneis. Hard minimum age: 56 Ma. 95% soft maximum age: 64 Ma. Prior setting: Lognormal distribution, mean= 0.78, St. Dev.= 0.8 (crown calibration). Calibration source: Near et al. 2012.

After time calibration the tree was annotated by adding heatmap for values of slope as calculated above (both number and size) by using webtool known as iPhylo (Li *et al.* 2025).

***Trait analysis***

Total number of TEs in each organism were calculated from the EDTA output file by counting each structurally (as described above) detected TE from the file. During this calculation the TE that do have exactly same start and end loci in same chromosome were considered as duplication and counted only once. We then calculated how many TEs are overlapping the genic area. Genic area included mRNA, rRNA, tRNA, LncRNA, snRNA, snoRNA, Pseudogenes, V gene region and C gene region. During this counting repetition of same TE as described above was taken care of by counting them once. We calculated TE in genic and Total TE for both number and Size. This was considered as an index of preference of TE to occupy genic region (trait 1). The relationship between TE (both number and size) and chromosome size for every organism is a significant straight line. The rate at which TE increases with increase in chromosome size was calculated from the slope of these straight-line relationships. These slopes were used as tolerance (trait 2) of a host to accumulate TEs. Evolution of these two traits across the above phylogenetic tree was analysed by reconstructing the ancestral trait reconstruction method. We have used R packages phytools (Revell 2024) and ape (Paradis and Schliep 2019.) to access methods like read.tree to read the phylogenetic time tree in Newick file format , read.table to read the trait file in txt format having name of the species in 1$^{st}$ column and trait values in the 2$^{nd}$ column (tab delimited 2 column txt file), fastAnc to re construct ancestral trait values for each ancestral nodes, contMap and setMap were used for plotting Continuous trait mapping with with the phylogeny having color gradient, phylosig was used with method = "K" (Blomberg, Garland and Ives 2003) to determine phylogenetic signals and fancyTree was used to plot phenogram95 (Revell 2013).

***Statistical analysis***

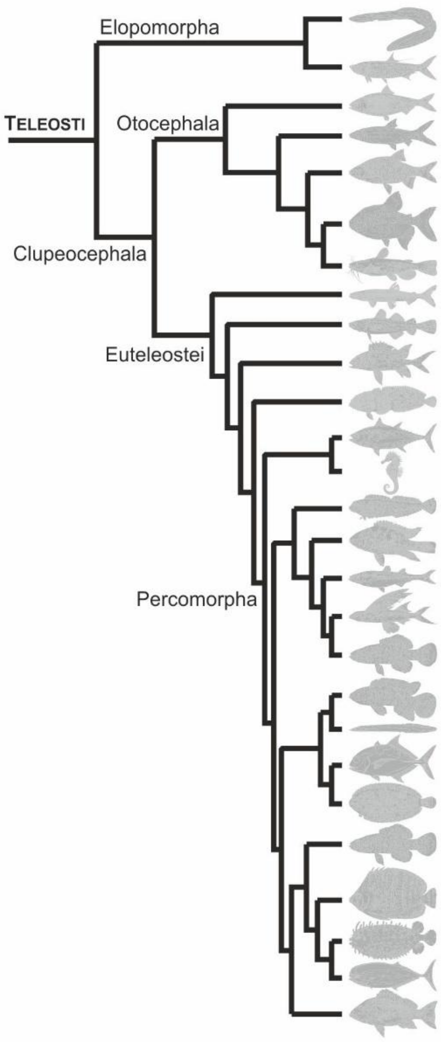


| Order | Families | Genera | Species |
|---|---|---|---|
| Anguilliformes | 1 | 1 | 1 |
| Elopiformes | 1 | 1 | 1 |
| Clupeiformes | 4 | 4 | 4 |
| Gonorynchiformes | 1 | 1 | 1 |
| Cypriniformes | 3 | 4 | 4 |
| Characiformes | 2 | 2 | 2 |
| Siluriformes | 3 | 3 | 3 |
| Osmeriformes | 1 | 1 | 1 |
| Gadiformes | 1 | 1 | 3 |
| Holocentriformes | 1 | 1 | 1 |
| Batrachoidiformes | 1 | 1 | 1 |
| Scombriformes | 1 | 2 | 3 |
| Syngnathiformes | 1 | 5 | 6 |
| Blenniiformes | 2 | 2 | 2 |
| Cichliformes | 1 | 5 | 6 |
| Atheriniformes | 1 | 1 | 1 |
| Beloniformes | 2 | 2 | 3 |
| Cyprinodontiformes | 3 | 4 | 5 |
| Anabantiformes | 1 | 1 | 1 |
| Synbranciformes | 1 | 1 | 1 |
| Carangoiformes | 2 | 2 | 2 |
| Pleuronectiformes | 2 | 5 | 7 |
| Centrarchiformes | 1 | 1 | 1 |
| Chetodontiformes | 1 | 1 | 1 |
| Tetraodontiformes | 1 | 1 | 1 |
| Spariformes | 1 | 2 | 2 |
| Perciformes | 6 | 9 | 12 |

**Fig. S1: Systematics of 78 Teleost fish genomes used for Transposable Element (TE) annotation and analysis.** The 78 species represent 27 out of total 40 (67.5%) Teleost orders.

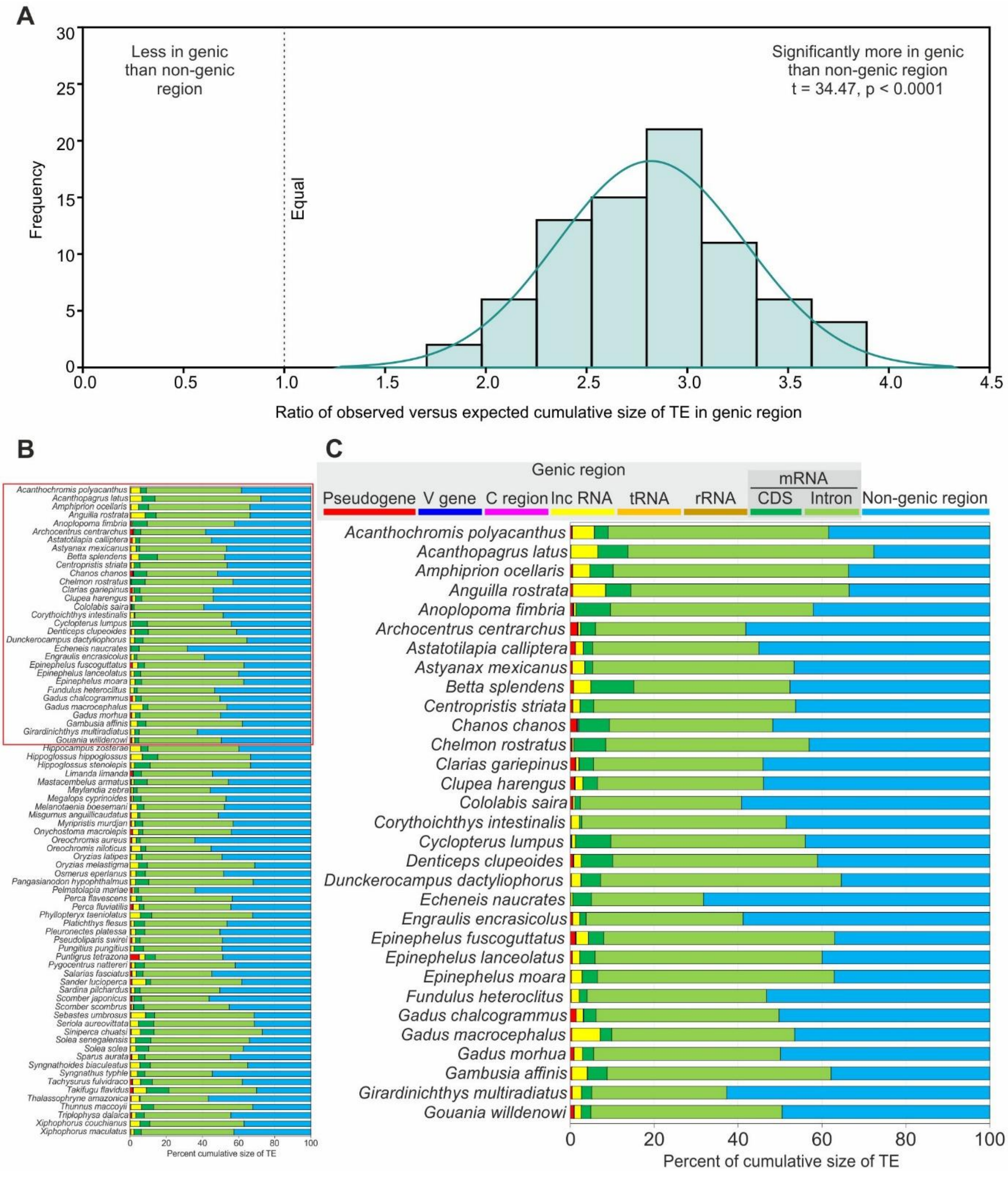


**Fig. S2. Distribution of TE in host genome based on size of TE (bp).** (A) Frequency of observed versus expected number of transposons in the genic region under random distribution. Significantly higher number of transposons in the genic region indicates preference of TE to genic region. (B) Distribution of TE in various genic region in 78 teleost fishes. (C) Selected view of (B) for higher resolution.

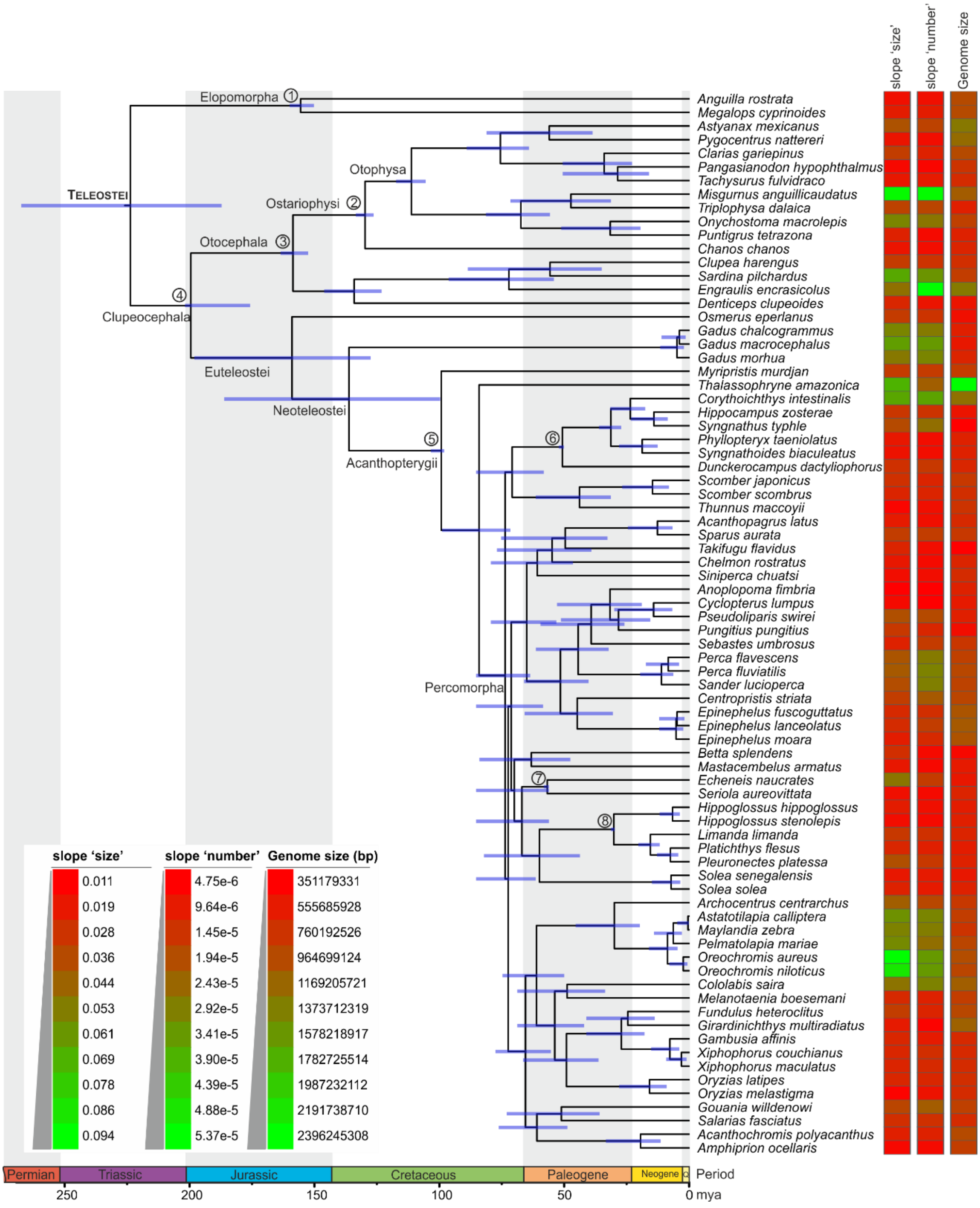


**Fig. S3: Time tree of 78 Teleost fish used for Transposable Element (TE) annotation and analysis.** RelTime analysis was performed on maximum likelihood tree obtained using best partition scheme and amino acid substitution model on 51 single copy genes. Slope of the relationship between size of the TE and chromosome size, slope of the relationship between TE numbers and chromosome size and the size of the genome are shown next to species labels.

### References


Benton, M.J., Donoghue, P.C., Asher, R.J., Friedman, M., Near, T.J. and Vinther, J., 2015. Constraints on the timescale of animal evolutionary history. https://doi.org/10.26879/424

Betancur-R, R., Broughton, R.E., Wiley, E.O., Carpenter, K., López, J.A., Li, C., Holcroft, N.I., Arcila, D., Sanciangco, M., Cureton Ii, J.C. and Zhang, F., 2013. The tree of life and a new classification of bony fishes. *PLoS currents*, *5*, pp.ecurrents-tol. 10.1371/currents.tol.53ba26640df0ccaee75bb165c8c26288

Blomberg, S.P., Garland Jr, T. and Ives, A.R., 2003. Testing for phylogenetic signal in comparative data: behavioral traits are more labile. *Evolution*, *57*(4), pp.717-745. https://doi.org/10.1111/j.0014-3820.2003.tb00285.xDigital Object Identifier (DOI)

Broughton, R.E., Betancur-R, R., Li, C., Arratia, G. and Ortí, G., 2013. Multi-locus phylogenetic analysis reveals the pattern and tempo of bony fish evolution. *PLoS currents*, *5*, pp.ecurrents-tol. 10.1371/currents.tol.2ca8041495ffafd0c92756e75247483e

Castresana, J., 2000. Selection of conserved blocks from multiple alignments for their use in phylogenetic analysis. *Molecular biology and evolution*, *17*(4), pp.540-552. https://doi.org/10.1093/oxfordjournals.molbev.a026334

Cooper, J.A. and Chapleau, F., 1998. Phylogenetic status of Paralichthodes algoensis (Pleuronectiformes: Paralichthodidae). *Copeia*, *1998*(2), pp.477-481. https://doi.org/10.2307/1447445

Edgar, R.C., 2004. MUSCLE: multiple sequence alignment with high accuracy and high throughput. *Nucleic acids research*, *32*(5), pp.1792-1797. https://doi.org/10.1093/nar/gkh340

Ellinghaus, D., Kurtz, S. and Willhoeft, U., 2008. LTRharvest, an efficient and flexible software for de novo detection of LTR retrotransposons. *BMC bioinformatics*, *9*(1), p.18. https://doi.org/10.1186/1471-2105-9-18

Ghezelayagh, A., Harrington, R.C., Burress, E.D., Campbell, M.A., Buckner, J.C., Chakrabarty, P., Glass, J.R., McCraney, W.T., Unmack, P.J., Thacker, C.E. and Alfaro, M.E., 2022. Prolonged morphological expansion of spiny-rayed fishes following the end-Cretaceous. *Nature Ecology & Evolution*, *6*(8), pp.1211-1220. https://doi.org/10.1038/s41559-022-01801-3

Grechishnikova, D. and Poptsova, M., 2016. Conserved 3′ UTR stem-loop structure in L1 and Alu transposons in human genome: possible role in retrotransposition. *BMC genomics*, *17*(1), p.992. https://doi.org/10.1186/s12864-016-3344-4

Han, J.S., 2010. Non-long terminal repeat (non-LTR) retrotransposons: mechanisms, recent developments, and unanswered questions. *Mobile Dna*, *1*(1), p.15. https://doi.org/10.1186/1759-8753-1-15

Harrington, R.C., Faircloth, B.C., Eytan, R.I., Smith, W.L., Near, T.J., Alfaro, M.E. and Friedman, M., 2016. Phylogenomic analysis of carangimorph fishes reveals flatfish asymmetry arose in a blink of the evolutionary eye. *BMC evolutionary biology*, *16*(1), p.224. https://doi.org/10.1186/s12862-016-0786-x



Harrington, R.C., Faircloth, B.C., Eytan, R.I., Smith, W.L., Near, T.J., Alfaro, M.E. and Friedman, M., 2016. Phylogenomic analysis of carangimorph fishes reveals flatfish asymmetry arose in a blink of the evolutionary eye. *BMC evolutionary biology*, *16*(1), p.224. https://doi.org/10.1186/s12862-016-0786-x

Harrington, R.C., Faircloth, B.C., Eytan, R.I., Smith, W.L., Near, T.J., Alfaro, M.E. and Friedman, M., 2016. Phylogenomic analysis of carangimorph fishes reveals flatfish asymmetry arose in a blink of the evolutionary eye. *BMC evolutionary biology*, *16*(1), p.224. https://doi.org/10.1186/s12862-016-0786-x

Hughes, L.C., Ortí, G., Huang, Y., Sun, Y., Baldwin, C.C., Thompson, A.W., Arcila, D., Betancur-R, R., Li, C., Becker, L. and Bellora, N., 2018. Comprehensive phylogeny of ray-finned fishes (Actinopterygii) based on transcriptomic and genomic data. *Proceedings of the National Academy of Sciences*, *115*(24), pp.6249-6254. https://doi.org/10.1073/pnas.1719358115

Kapitonov, V.V. and Jurka, J., 2006. Self-synthesizing DNA transposons in eukaryotes. *Proceedings of the National Academy of Sciences*, *103*(12), pp.4540-4545. https://doi.org/10.1073/pnas.0600833103

Kapitonov, V.V. and Jurka, J., 2007. Helitrons on a roll: eukaryotic rolling-circle transposons. *TRENDS in Genetics*, *23*(10), pp.521-529. 10.1016/j.tig.2007.08.004

Li, Y., Peng, C., Chi, F., Huang, Z., Yuan, M., Zhou, X. and Jiang, C., 2025. The iPhylo suite: an interactive platform for building and annotating biological and chemical taxonomic trees. *Briefings in Bioinformatics*, *26*(1), p.bbae679. https://doi.org/10.1093/bib/bbae679

Near, T.J., Eytan, R.I., Dornburg, A., Kuhn, K.L., Moore, J.A., Davis, M.P., Wainwright, P.C., Friedman, M. and Smith, W.L., 2012. Resolution of ray-finned fish phylogeny and timing of diversification. *Proceedings of the National Academy of Sciences*, *109*(34), pp.13698-13703. https://doi.org/10.1073/pnas.1206625109

Nguyen, L.T., Schmidt, H.A., Von Haeseler, A. and Minh, B.Q., 2015. IQ-TREE: a fast and effective stochastic algorithm for estimating maximum-likelihood phylogenies. *Molecular biology and evolution*, *32*(1), pp.268-274. https://doi.org/10.1093/molbev/msu300

Ou S, Su W, Liao Y, Chougule K, Agda JR, Hellinga AJ, Lugo CSB, Elliott TA, Ware D, Peterson T and Jiang N (2019). Benchmarking transposable element annotation methods for creation of a streamlined, comprehensive pipeline. *Genome biology*, 20:1-18. https://doi.org/10.1186/s13059-019-1905-y

Ou, S. and Jiang, N., 2018. LTR_retriever: a highly accurate and sensitive program for identification of long terminal repeat retrotransposons. *Plant physiology*, *176*(2), pp.1410-1422. https://doi.org/10.1104/pp.17.01310

Paradis, E. and Schliep, K., 2019. ape 5.0: an environment for modern phylogenetics and evolutionary analyses in R. *Bioinformatics*, *35*(3), pp.526-528. https://doi.org/10.1093/bioinformatics/bty633

Revell, L.J., 2013. Two new graphical methods for mapping trait evolution on phylogenies. *Methods in Ecology and Evolution*, 4(8), pp.754-759. https://doi.org/10.1111/2041-210X.12066

Revell, L.J., 2024. phytools 2.0: an updated R ecosystem for phylogenetic comparative methods (and other things). *PeerJ*, *12*, p.e16505. https://doi.org/10.7717/peerj.16505

Sakamoto, K., Uyeno, T. and Micklich, N., 2004. Oligopleuronectes germanicus gen. et sp. nov., an Oligocene pleuronectid flatfish from Frauenweiler, S-Germany. *Bulletin of the National Science Museum, Tokyo, Series C*, *30*, pp.89-94.

Shi, J. and Liang, C., 2019. Generic repeat finder: a high-sensitivity tool for genome-wide de novo repeat detection. *Plant physiology*, *180*(4), pp.1803-1815. https://doi.org/10.1104/pp.19.00386

Smit, A.F.A., Hubley, R. and Green, P., 2015. RepeatModeler Open-1.0. 2008–2015. *Seattle, USA: Institute for Systems Biology*, *1*, p.2018. www.repeatmasker.Org 10.1073/pnas.1921046117

Smit, A.F.A., Hubley, R. and Green, P., 2015. *RepeatMasker Open-4.0. 2013–2015* [online] www.repeatmasker.org

Su, W., Gu, X. and Peterson, T., 2019. TIR-Learner, a new ensemble method for TIR transposable element annotation, provides evidence for abundant new transposable elements in the maize genome. *Molecular plant*, *12*(3), pp.447-460. 10.1016/j.molp.2019.02.008

Tamura, K., Dudley, J., Nei, M. and Kumar, S., 2007. MEGA4: molecular evolutionary genetics analysis (MEGA) software version 4.0. *Molecular biology and evolution*, *24*(8), pp.1596-1599. https://doi.org/10.1093/molbev/msm092

Thomas, J. and Pritham, E.J., 2015. Helitrons, the eukaryotic rolling-circle transposable elements. *Mobile DNA III*, pp.891-924. https://doi.org/10.1128/9781555819217.ch40Digital Object Identifier (DOI)

Waterhouse, R.M., Seppey, M., Simão, F.A., Manni, M., Ioannidis, P., Klioutchnikov, G., Kriventseva, E.V. and Zdobnov, E.M., 2018. BUSCO applications from quality assessments to gene prediction and phylogenomics. *Molecular biology and evolution*, *35*(3), pp.543-548. https://doi.org/10.1093/molbev/msx319

Xiong, W., He, L., Lai, J., Dooner, H.K. and Du, C., 2014. HelitronScanner uncovers a large overlooked cache of Helitron transposons in many plant genomes. *Proceedings of the National Academy of Sciences*, *111*(28), pp.10263-10268. https://doi.org/10.1073/pnas.1410068111

Xu, L., Zhang, Y., Su, Y., Liu, L., Yang, J., Zhu, Y. and Li, C., 2010. Structure and evolution of full-length LTR retrotransposons in rice genome. *Plant systematics and evolution,* 287(1), pp.19-28. https://doi.org/10.1007/s00606-010-0285-2

Xu, Z. and Wang, H., 2007. LTR_FINDER: an efficient tool for the prediction of full-length LTR retrotransposons. *Nucleic acids research*, *35*(suppl_2), pp.W265-W268. https://doi.org/10.1093/nar/gkm286
